# Parietal representations of stimulus features are amplified during memory retrieval and flexibly aligned with top-down goals

**DOI:** 10.1101/280974

**Authors:** Serra E. Favila, Rosalie Samide, Sarah C. Sweigart, Brice A. Kuhl

## Abstract

In studies of human episodic memory, the phenomenon of reactivation has traditionally been observed in regions of occipitotemporal cortex (OTC) involved in sensory experience. However, reactivation also occurs in lateral parietal cortex (LPC), and recent evidence indicates that reactivation of stimulus-specific information may be stronger in LPC than in OTC. These observations raise a number of questions about the nature of memory representations in LPC and their relation to representations in OTC. Here, we report two fMRI experiments that quantify stimulus feature information (color and object category) within LPC and OTC, separately during perception and memory retrieval, in male and female human subjects. Across both experiments, we show a clear dissociation between OTC and LPC: while feature information in OTC is relatively stronger during perception than memory, feature information in LPC is relatively stronger during memory than perception. Thus, while OTC and LPC represent common stimulus features, they preferentially represent this information during different stages. We show that this transformation of feature information across regions co-occurs with stimulus-level reinstatement within LPC and high-level OTC. In Experiment 2, we consider whether feature information in LPC during memory retrieval is flexibly and dynamically shaped by top-down goals. Indeed, we find that dorsal LPC preferentially represents retrieved feature information that addresses current goals. In contrast, ventral LPC represents retrieved features independent of current goals. Collectively, these findings provide insight into the nature and significance of mnemonic representations in LPC and constitute an important bridge between putative mnemonic and control functions of parietal cortex.

## Introduction

Traditional models of episodic memory posit that sensory activity evoked during perception is reactivated during recollection (Kosslyn, 1980; Damasio, 1989). There is considerable evidence for such reactivation in visual regions of occipitotemporal cortex (OTC) (Wheeler et al., 2000; Polyn et al., 2005). However, recent human neuroimaging work indicates that stimulus information is also reactivated in lateral parietal cortex (LPC) (Kuhl and Chun, 2014; Chen et al., 2016; Lee and Kuhl, 2016; Xiao et al., 2017). While these findings accord with evidence for univariate increases in LPC BOLD activity during successful remembering (Wagner et al., 2005; Kuhl and Chun, 2014), they also raise important questions about whether and how representations of retrieved memories differ between LPC and OTC.

Univariate fMRI studies have consistently found that, in contrast to sensory regions, ventral LPC exhibits low activation when perceptual events are experienced but high activation when these events are successfully retrieved (Daselaar, 2009; Kim et al., 2010). The idea that LPC is relatively more involved in memory retrieval than perception has also received support from recent pattern-based fMRI studies. Long, Lee, and Kuhl (2016) found that reactivation of previously learned visual category information was stronger in the default mode network (which includes ventral LPC) than in OTC (see also Chen et al., 2016), whereas the reverse was true of category information during perception. Similarly, Xiao and colleagues (2017) found that stimulus-specific representations of retrieved stimuli were relatively stronger in LPC than in high-level visual areas, whereas stimulus-specific representations of perceived stimuli showed the opposite pattern.

Collectively, these studies raise the intriguing idea that reactivation–defined as the consistency of activation patterns across perception and retrieval–may not fully capture how memories are represented during recollection. Rather, there may be a transformation of information from sensory regions during perception to higher-order regions (including LPC) during retrieval. Critically, however, previous studies have not measured or compared representations of stimulus features in OTC and LPC during perception vs. memory. This leaves open the important question of whether the same features represented in OTC during perception are also represented in LPC during retrieval, or whether these two regions represent different aspects of the stimuli across processing stages (Xiao et al., 2017). In addition, there is currently little evidence addressing how stimulus-level representations in LPC relate to, or emerge from, feature-level representations (Kuhl and Chun, 2014). Finally, when considering the proposed attentional and control functions of parietal cortex (Behrmann, 2004), consideration of feature-level representations in LPC is particularly important because LPC may play a role in flexibly aligning retrieved representations with behavioral goals (Kuhl et al., 2013). By this account, representations in LPC may be biased toward the specific stimulus features that are relevant to current memory demands.

We conducted two fMRI experiments designed to directly compare visual stimulus representations during perception and memory in OTC and LPC. Stimuli were images of common objects with two visual features of interest: color and object category (Fig. 1). In both experiments (Fig. 2*A*), human subjects learned word-image associations prior to a scan session. During scanning, subjects completed separate perception and memory tasks (Fig. 2*B*). During perception trials, subjects viewed the image stimuli. During memory trials, subjects were presented with word cues and recalled the associated images. The key difference between Experiments 1 and 2 occurred during memory trials. In Experiment 1, subjects retrieved each image as vividly as possible, whereas in Experiment 2 subjects alternately recalled only the color or object feature of each image as vividly as possible. Using data from both experiments, we evaluated the relative strength of color and object feature information in OTC and LPC during stimulus perception and memory. We also compared the strength of feature-level and stimulus-level reinstatement in these regions. Using data from Experiment 2, we evaluated the role of top-down goals on mnemonic feature representations, specifically testing for differences in goal-sensitivity across LPC subregions (Sestieri et al., 2017).

**Figure 1.**
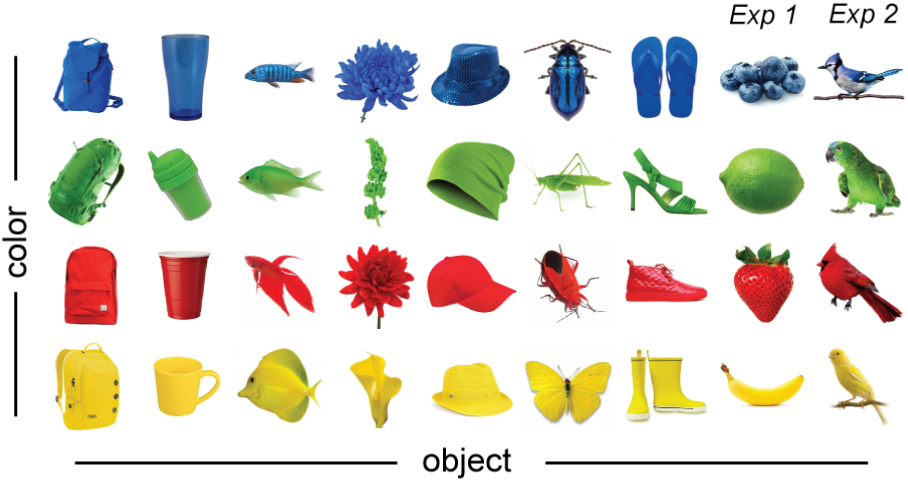
Stimuli. In both experiments, stimuli were images of 32 common objects. Each object was a unique conjunction of one of four color features and one of eight object features. Colors were blue, green, red, and yellow. Object features were backpacks, cups, fish, flowers, hats, insects, shoes, fruit (Experiment 1 only), and birds (Experiment 2 only). See also Materials and Methods.

**Figure 2.**
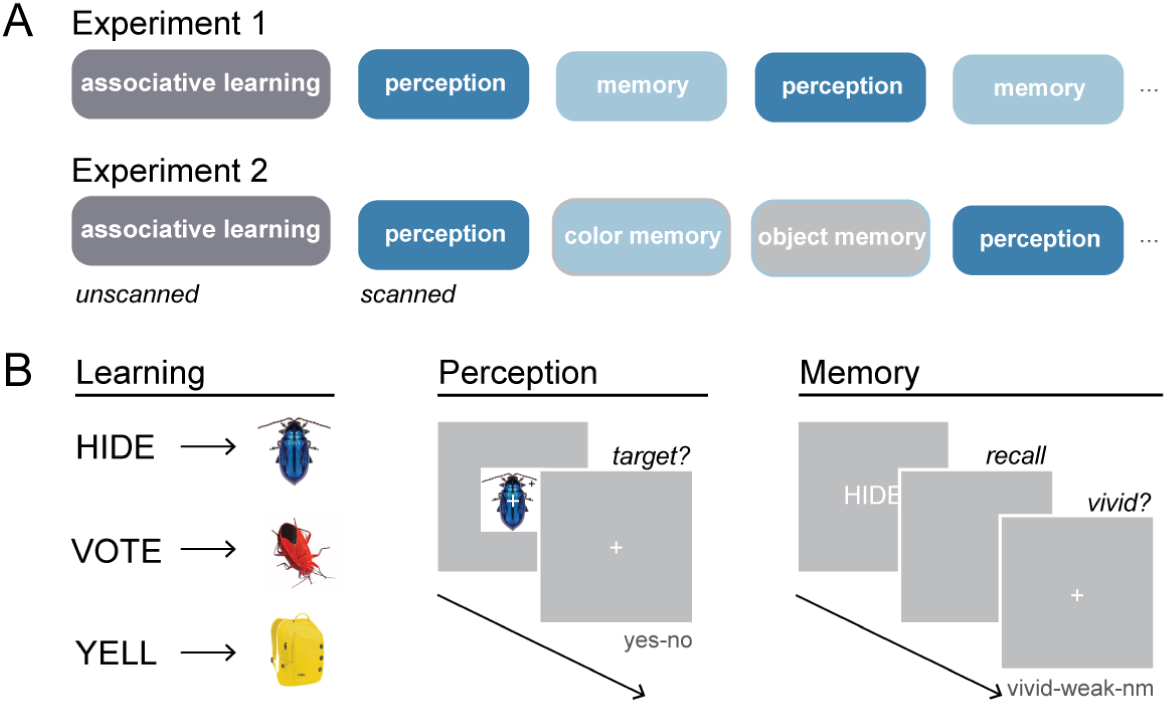
Experimental design and task structure. ***A***, In both experiments, human subjects learned word-image paired associates prior to scanning. In the scanner, subjects viewed and recalled the image stimuli in alternate perception and memory runs. In Experiment 2, subjects performed two goal-dependent memory tasks, during which they were cued to selectively recall only the color feature or only the object feature of the associated image. ***B***, Subjects learned 32 word-image pairs to a 100% criterion in the behavioral training session. During scanned perception trials, subjects were briefly presented with a stimulus. Subjects judged whether a small infrequent visual target was present or absent on the stimulus. During scanned memory trials, subjects were presented with a previously studied word cue, and recalled the associated stimulus (Experiment 1) or only the color or object feature of the associated stimulus (Experiment 2). After a brief recall period, subjects made a vividness judgment about the quality of their recollection (vivid, weak, no memory). See also Materials and Methods.

## Material and Methods

### Subjects

Forty-seven male and female human subjects were recruited from the New York University (Experiment 1) and University of Oregon (Experiment 2) communities. All subjects were right-handed native English speakers between the ages of 18 and 35 who reported normal or corrected-to-normal visual acuity, normal color vision, and no history of neurological or psychiatric disorders. Subjects participated in the study after giving written informed consent to procedures approved by the New York University or University of Oregon Institutional Review Boards. Of the 24 subjects recruited for Experiment 1, seven subjects were excluded from data analysis due to poor data quality owing to excessive head motion (n = 3), sleepiness during the scan (n = 2), or poor performance during memory scans (n = 2, < 75% combined vivid memory and weak memory responses). This yielded a final data set of 17 subjects for Experiment 1 (19 - 31 years old, 7 males). Of the 23 subjects recruited for Experiment 2, two subjects withdrew from the study prior to completion due to either a scanner error (n = 1) or discomfort during the scan (n = 1). An additional four subjects were excluded from data analysis due to: an abnormality detected in the acquired images (n = 1), poor data quality owing to excessive head motion (n = 2), or poor performance during memory scans (n = 1, < 75% combined vivid memory and weak memory responses). This yielded a final data set of 17 subjects for Experiment 2 (18 - 31 years old, 8 males).

### Stimuli

Stimuli for Experiment 1 consisted of 32 unique object images (Fig. 1). Each object was a unique conjunction of one of four color features (blue, green, red, yellow) and one of eight object features (backpacks, cups, fish, flowers, fruit, hats, insects, shoes). Thirty-two closely matched foil images were also used in the behavioral learning session. Stimuli for Experiment 2 were identical to those from Experiment 1, with the exception that the object category fruits was replaced with the category birds. All images were 225 x 225 pixels, with the object rendered on a white background. Word cues consisted of 32 common verbs and were the same for both experiments.

### Tasks and procedure

#### Experiment 1

The experiment began with a behavioral session, during which subjects learned 32 unique word-image associations to 100% criterion. A scan session immediately followed completion of the behavioral session. During the scan, subjects participated in two types of runs: 1) perception, where they viewed the object images without the corresponding word cues and 2) memory, where they were presented with the word cues and recalled the associated object images (Fig. 2*A, B*). Details for each of these phases are described below.

Immediately prior to scanning, subjects learned 32 word-image associations through interleaved study and test blocks. For each subject, the 32 word cues were randomly assigned to each of 32 object images. During study blocks, subjects were presented with the 32 word-image associations in random order. On a given study trial, the word cue was presented for 2 s, followed by the associated object image for 2 s. A fixation cross was presented centrally for 2 s before the start of the next trial. Subjects were instructed to learn the associations in preparation for a memory test, but no responses were required. During test blocks, subjects were presented with the 32 word cues in random order and tested on their memory for the associated image. On each test trial, the word cue was presented for .5 s and was followed by a blank screen for 3.5 s, during which subjects were instructed to try to recall the associated image as vividly as possible for the entire 3.5 s. After this period elapsed, a test image was presented. The test image was either the correct object (target), an object that had been associated with a different word cue (old), or a novel object that was highly similar (same color and object category) to the target (lure). These trial types occurred with equal probability. For each test image, subjects had up to 5 s to make a yes/no response indicating whether or not the test image was the correct associate. After making a response, subjects were shown the target image for 1 s as feedback. After feedback, a fixation cross was presented centrally for 2 s before the start of the next trial. Lure trials were included to ensure that subjects formed detailed memories of each object (i.e., that subjects could discriminate between the target image and another image with the same combination of features). Subjects alternated between study and test blocks until they completed a minimum of 6 blocks of each type and achieved 100% accuracy on the test.

Once in the scanner, subjects participated in two types of runs: perception and memory. During perception runs, subjects viewed the object images one at a time while performing a cover task of detecting black crosses that appeared infrequently on images. On a given perception trial, the object image was overlaid with a central white fixation cross and presented centrally on a gray background for .5 s. The central white fixation cross was then presented alone on a gray background for 3.5 s before the start of the next trial. Subject were instructed to maintain fixation on the central fixation cross and monitor for a black cross that appeared at a random location within the borders of a randomly selected 12.5% of images. Subjects were instructed to judge whether a target was present or absent on the image and indicate their response with a button press. Each perception run consisted of 32 object perception trials (1 trial per stimulus) and 8 null fixation trials in random order. Null trials consisted of a central white fixation cross on a gray background presented for 4 s and were randomly interleaved with the object trials thereby creating jitter. Every run also contained 8 s of null lead in and 8 s of null lead out time during which a central white fixation cross on a gray background was presented.

During memory runs, subjects were presented with the word cues one at a time, recalled the associated images, and evaluated the vividness of their recollections. On each memory trial, the word cue was presented centrally in white characters on a gray background for .5 s. This was followed by a 2.5 s recall period where the screen was blank. Subjects were instructed to use this period to recall the associated image from memory and to hold it in mind as vividly as possible for the entire duration of the blank screen. At the end of the recall period, a white question mark on a gray background was presented for 1 s, prompting subjects to make one of three memory vividness responses via button box: “vividly remembered”, “weakly remembered”, “not remembered”. The question mark was replaced by a central white fixation cross, which was presented for 2 s before the start of the next trial. Responses were recorded if they were made during the question mark or the ensuing fixation cross. As in perception runs, each memory run consisted of 32 object memory trials (1 trial per stimulus) and 8 null fixation trials in random order. Null trials consisted of a central white fixation cross on a gray background presented for 6 s, and as in perception runs, provided jitter. Each run contained 8 s of null lead in and 8 s of null lead out time during which a central white fixation cross on a gray background was presented.

For both perception and memory tasks, trial orders were randomly generated for each subject and run. Subjects alternated between perception and memory runs, performing as many runs of each task as could be completed during the scan session (range = 7-10, mean = 8.41). Thus, there were between 7 and 10 repetitions of each stimulus across all perception trials and 7 to 10 repetitions of each stimulus across all memory trials. All stimuli were displayed on a projector at the back of the scanner bore, which subjects viewed through a mirror attached to the head coil. Subjects made responses for both tasks on an MR-compatible button box.

#### Experiment 2

As in Experiment 1, Experiment 2 began with a behavioral session, during which subjects learned 32 unique word-image associations to 100% criterion. A scan session immediately followed. During the scan, subjects participated in both perception and memory runs. In contrast to Experiment 1, subjects performed one of two goal-dependent memory tasks during memory runs: 1) color memory, where they selectively recalled the color feature of the associated image from the word cue; 2) object memory, where they selectively recalled the object feature of the associated image from the word cue (Fig. 2*A, B*). Details of each phase of the experiment, in relation to Experiment 1, are described below.

Subjects learned 32 word-image associations following the same procedure as in Experiment 1. Once in the scanner, subjects participated in three types of runs: perception, color memory, and object memory. Procedures were the same as in Experiment 1 unless noted. During perception runs, subjects viewed the object images one at a time while performing a cover task of detecting black crosses that infrequently appeared on images. On a given perception trial, the object image was overlaid with a central white fixation cross and presented centrally on a gray background for .75 s. The central white fixation cross was then presented alone on a gray background for either 1.25, 3.25, 5.25, 7.25, or 9.25 s (25%, 37.5%, 18.75%, 12.5%, 6.25% of trials per run, respectively) before the start of the next trial. These interstimulus intervals were randomly assigned to trials. Subjects performed the detection task as in Experiment 1. Each perception run consisted of 64 perception trials (2 trials per stimulus) in random order, with lead in and lead out time as in Experiment 1.

During color and object memory runs, subjects were presented with the word cues one at a time, recalled either the color or object feature of the associated images, and evaluated the vividness of their recollections. On each memory trial, the word cue was presented centrally in white characters on a gray background for .3 s. This was followed by a 2.2 s recall period where the screen was blank. Subjects were instructed to use this period to recall only the cued feature of the associated image from memory and to hold it in mind as vividly as possible for the entire duration of the blank screen. At the end of the recall period, a white fixation cross was presented centrally on a gray background for either 1.5, 3.5, 5.5, 7.5, or 9.5 s (37.5%, 25%, 18.75%, 12.5%, 6.25% of trials per run, respectively), prompting subjects to make one of three memory vividness decisions via button box as in Experiment 1. The interstimulus intervals were randomly assigned to trials. Color and object memory runs consisted of 64 memory trials (2 trials per stimulus) presented in random order, with lead in and lead out time as in Experiment 1

All subjects completed 4 perception runs, 4 color memory runs, and 4 object memory runs, with each stimulus presented twice in every run. Thus, there were 8 repetitions of each stimulus for each run type. Runs were presented in four sequential triplets, with each triplet composed of one perception run followed by color and object memory runs in random order. As in Experiment 1, stimuli were displayed on a projector at the back of the scanner bore, which subjects viewed through a mirror attached to the head coil. Subjects made responses for all three tasks on an MR-compatible button box.

### MRI acquisition

#### Experiment 1

Images were acquired on a 3T Siemens Allegra head-only MRI system at the Center for Brain Imaging at New York University. Functional data were acquired with a T2*-weighted echo-planar imaging (EPI) sequence with partial coverage (repetition time = 2 s, echo time = 30 ms, flip angle = 82°, 34 slices, 2.5 x 2.5 x 2.5 mm voxels) and an 8 channel occipital surface coil. Slightly oblique coronal slices were aligned approximately 120° with respect to the calcarine sulcus at the occipital pole and extended anteriorly covering the occipital lobe, ventral temporal cortex and posterior parietal cortex. A whole-brain T1-weighted magnetization-prepared rapid acquisition gradient echo (MPRAGE) 3D anatomical volume (1x 1 x 1 mm voxels) was also collected.

#### Experiment 2

Images were acquired on a 3T Siemens Skyra MRI system at the Robert and Beverly Lewis Center for NeuroImaging at the University of Oregon. Functional data were acquired using a T2*-weighted multiband EPI sequence with whole-brain coverage (repetition time = 2 s, echo time = 25 ms, flip angle = 90°, multiband acceleration factor = 3, inplane acceleration factor = 2, 72 slices, 2 x 2x 2 mm voxels) and a 32 channel head coil. Oblique axial slices were aligned parallel to the plane defined by the anterior and posterior commissures. A whole-brain T1-weighted MPRAGE 3D anatomical volume (1 x 1 x 1 mm voxels) was also collected.

### fMRI processing

FSL v5.0 (Smith et al., 2004) was used for functional image preprocessing. The first four volumes of each functional run were discarded to allow for T1 stabilization. To correct for head motion, each run’s timeseries was realigned to its middle volume. Each timeseries was spatially smoothed using a 4 mm full width at half maximum Gaussian kernel and high-pass filtered using Gaussian-weighted least squares straight line fitting with *σ* = 64.0 s. Volumes with motion relative to the previous volume greater than 1.25 mm in Experiment 1 (half the width of a voxel) or greater than .5 mm in Experiment 2 were excluded from subsequent analyses. A lower threshold was chosen for Experiment 2 due to high motion artifact susceptibility in multiband sequences. Freesurfer v5.3 (Fischl, 2012) was used to perform segmentation and cortical surface reconstruction on each subject’s anatomical volume. Boundary-based registration was used to compute the alignment between each subject’s functional data and their anatomical volume.

All fMRI processing was performed in individual subject space. To estimate the neural pattern of activity evoked by the perception and memory of every stimulus, we conducted separate voxelwise general linear model (GLM) analyses of each subject’s smoothed timeseries data from the perception and memory runs in each experiment. Perception models included 32 regressors of interest corresponding to the presentation of each stimulus. Events within these regressors were constructed as boxcars with stimulus presentation duration convolved with a canonical double-gamma hemodynamic response function. Six realignment parameters were included as nuisance regressors to control for motion confounds. First-level models were estimated for each run using Gaussian least squares with local autocorrelation correction (“prewhitening”). Parameter estimates and variances for each regressor were then registered into the space of the first run and entered into a second-level fixed effects model. This produced *t*-maps representing the activation elicited by by viewing each stimulus for each subject. No normalization to a group template was performed. Memory models were estimated using the same procedure, with a regressor of interest corresponding to the recollection of each of the 32 stimuli. All recollection events were constructed as boxcars with a combined cue plus recall duration before convolution. This produced *t*-maps representing the activation elicited by remembering each stimulus relative to baseline for each subject. For Experiment 2, three memory models were estimated: one that included all memory trials from both tasks, and two task-specific models that included only color memory trials or only object memory trials.

### Region of interest definition

ROIs (Fig. 3) were produced for each subject in native subject space using multiple group-defined atlases. Our choice of group atlas for each broader cortical region of interest was based on our assessment of the best validated method for parcellating regions in that area. For retinotopic regions in OTC, we relied on a probabilistic atlas published by Wang et al. (2014). We combined bilateral V1v and V1d regions from this atlas to produce a V1 ROI and bilateral LO1 and LO2 regions to produce an LO ROI. For high-level OTC, we used the output of Freesurfer segmentation routines to combine bilateral fusiform gyrus, collateral sulcus, and lateral occipitemporal sulcus cortical labels to create a ventral temporal cortex (VTC) ROI. To subdivide LPC, we first selected the lateral parietal nodes of networks 5, 12, 13, 15, 16, and 17 of the 17 network resting state atlas published by Yeo et al. (2011). We refer to parietal nodes from Network 12 and 13 (subcomponents of the frontoparietal control network) as dorsal lateral intraparietal sulcus (dLatIPS) and ventral lateral intraparietal sulcus (vLatIPS), respectively. We altered the parietal node of Network 5 (dorsal attention network) by eliminating vertices in lateral occipital cortex and by subdividing it along the intraparietal sulcus into a dorsal region we refer to as posterior intraparietal sulcus (pIPS) and an ventral region we call ventral IPS (vIPS), following (Sestieri et al., 2017). The ventral region also corresponds closely to what others have called PGp (Caspers et al., 2012; Glasser et al., 2016). Finally, due to their small size, we combined the parietal nodes of Networks 15, 16, and 17 (subcomponents of the default mode network) into a region we collectively refer to as angular gyrus (AnG). All regions were first defined on Freesurfer’s average cortical surface (shown in Fig. 3) and then reverse-normalized to each subject’s native anatomical surface. They were then projected into the volume at the resolution of the functional data to produce binary masks.

**Figure 3.**
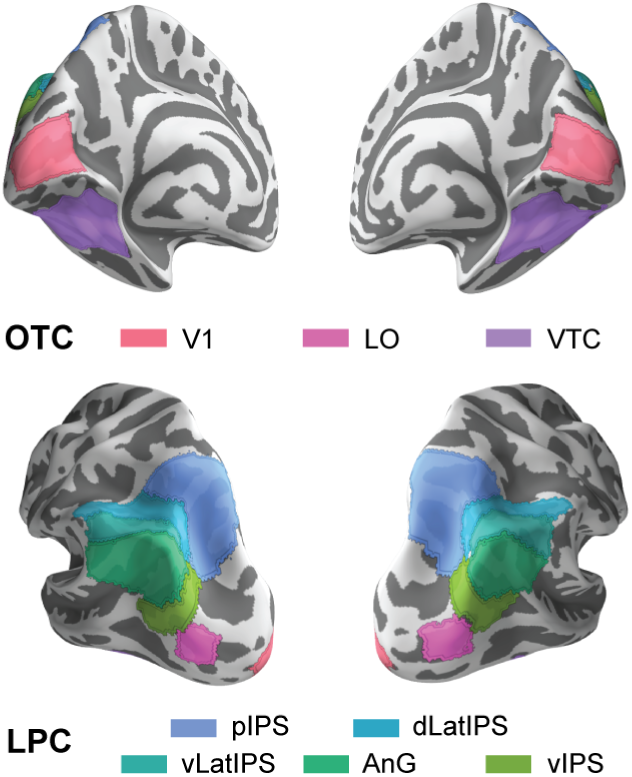
Regions of interest. Anatomical ROIs visualized on the Freesurfer average cortical surface. OTC ROIs included V1 and LO, defined using a group atlas of retinotopic regions (Wang et al., 2014), and VTC, defined using Freesurfer segmentation protocols. LPC ROIs included 5 ROIs that spanned dorsal and ventral LPC: pIPS, dLatIPS, vLatIPS, AnG, and vIPS. These ROIs were based on a group parcellation of cortical regions estimated from spontaneous activity (Yeo et al., 2011). All ROIs were transformed to subjects’ native anatomical surfaces and then into functional volume space prior to analysis. See also Materials and Methods.

### Experimental design and statistical analysis

Our experimental design for Experiment 1 included two types of cognitive tasks, which subjects performed in different fMRI runs–perception of visual stimuli, and retrieval of the same stimuli from long-term memory. Each of the 32 stimuli had one of four color features and one of eight object features. Experiment 2 was performed on an independent sample of subjects, and had a similar design to Experiment 1, except that subjects in Experiment 2 performed two goal-dependent versions of the memory retrieval task: color memory and object memory (see Tasks and Procedure). Our sample size for each experiment was consistent with similar fMRI studies in the field and was determined before data collection. Our dependent variables of interest for both experiments were stimulus-evoked BOLD activity patterns. In each experiment, separate *t*-maps were obtained for each stimulus from the perception and memory runs (see fMRI processing). Experiment 2 memory *t*-maps were derived from a single model that collapsed across the two goal-dependent memory tasks except when testing for goal-related effects. When testing for goal-related effects, we used *t*-maps that were separately estimated from the color and object memory tasks. We intersected all *t*-maps with binary ROI masks to produce stimulus-evoked voxel patterns for each ROI. Our ROIs included early and high-level visual areas in OTC that we believed would be responsive to the features of our stimuli, as well as regions spanning all of LPC (see Regions of interest definition). Analyses focused on cortical regions at multiple levels of spatial granularity. In order to evaluate whether perceptually-based and memory-based processing differed between LPC and OTC, we grouped data from individual ROIs according to this distinction and evaluated effects of ROI group (OTC, LPC). Given prior work implicating dorsal parietal cortex in top-down attention (Corbetta and Shulman, 2002), we also tested for differences in goal-modulated memory processing between dorsal and ventral LPC regions. To do this, we grouped data from individual LPC ROIs along a well-accepted boundary (the intraparietal sulcus) and evaluated effects of LPC subregion (dorsal, ventral). We report follow-up statistical tests performed on data from individual ROIs in Tables 1-3. All statistical tests performed on BOLD activity patterns (described below) were implemented in R v3.4. All *t*-tests were two-tailed, and with the exception of tests performed at the individual ROI level, all tests were assessed at alpha = 0.05. Tests in the 8 individual ROIs are reported in tables. Here, uncorrected *p*-values are reported with significance after correcting for multiple comparisons indicated. A conservative Bonferroni-corrected *p*-value of 0.05/8 = 0.00625 was used.

**Table 1.** Feature information during perception and memory in individual ROIs

| ROI | Perception |  |  |  | Memory |  |  |  |
| --- | --- | --- | --- | --- | --- | --- | --- | --- |
|  | Color |  | Object |  | Color |  | Object |  |
| | $t_{33}$ | $p$ | $t_{33}$ | $p$ | $t_{33}$ | $p$ | $t_{33}$ | $p$ |
| V1 | 2.32 | 0.027 | 3.29 | 0.002* | 2.55 | 0.015 | 1.18 | 0.246 |
| LO | 0.83 | 0.410 | 5.04 | <0.001* | -0.41 | 0.687 | 3.58 | 0.001* |
| VTC | -0.97 | 0.338 | 5.00 | <0.001* | 0.87 | 0.390 | 2.54 | 0.016 |
| pIPS | -2.24 | 0.032 | 0.18 | 0.858 | 1.82 | 0.078 | 2.72 | 0.010 |
| dLatIPS | -2.81 | 0.008 | 0.18 | 0.855 | 0.64 | 0.528 | 2.39 | 0.023 |
| vLatIPS | -1.81 | 0.080 | 0.66 | 0.513 | 1.76 | 0.087 | 3.15 | 0.003* |
| AnG | 0.10 | 0.919 | 0.36 | 0.718 | 3.48 | 0.001* | 3.48 | 0.001* |
| vIPS | 0.31 | 0.761 | 2.82 | 0.008 | 2.18 | 0.036 | 3.48 | 0.001* |

One sample $t$ -tests comparing feature information to chance (zero).
\* $p < 0.00625$ following multiple comparisons correction for 8 ROIs

**Table 2.** Feature and stimulus reinstatement in individual ROIs

| ROI | Color |  | Object |  | Stimulus > Color + Object |  |
| --- | --- | --- | --- | --- | --- | --- |
| | $t_{33}$ | $p$ | $t_{33}$ | $p$ | $t_{33}$ | $p$ |
| V1 | 0.56 | 0.582 | 1.16 | 0.253 | 0.42 | 0.674 |
| LO | 3.11 | 0.004* | 2.27 | 0.030 | -0.87 | 0.389 |
| VTC | 0.53 | 0.597 | 2.10 | 0.044 | 2.30 | 0.028 |
| pIPS | 1.29 | 0.207 | 0.60 | 0.556 | 2.47 | 0.019 |
| dLatIPS | 2.10 | 0.043 | -0.64 | 0.524 | 1.61 | 0.118 |
| vLatIPS | 2.04 | 0.050 | -0.59 | 0.560 | 1.92 | 0.063 |
| AnG | 1.94 | 0.062 | 0.75 | 0.461 | 0.65 | 0.519 |
| vIPS | 1.20 | 0.239 | 0.91 | 0.368 | 3.12 | 0.004* |

One sample $t$ -tests comparing feature reinstatement to chance (zero) and paired sample $t$ -tests comparing stimulus reinstatement to summed feature reinstatement.
\* $p < 0.00625$ following multiple comparisons correction for 8 ROIs

**Table 3.** Feature information during memory by goal-relevance in individual ROIs

| ROI | Relevant |  | Irrelevant |  | Relevant > Irrelevant |  |
| --- | --- | --- | --- | --- | --- | --- |
| | $t_{16}$ | $p$ | $t_{16}$ | $p$ | $t_{16}$ | $p$ |
| V1 | -1.11 | 0.285 | 2.24 | 0.040 | -2.10 | 0.052 |
| LO | -0.28 | 0.780 | 0.53 | 0.602 | -0.58 | 0.568 |
| VTC | 0.54 | 0.595 | 0.99 | 0.336 | -0.31 | 0.759 |
| pIPS | 1.85 | 0.084 | -0.06 | 0.953 | 1.79 | 0.092 |
| dLatIPS | 1.80 | 0.092 | -0.76 | 0.458 | 2.38 | 0.030 |
| vLatIPS | 3.53 | 0.003* | 1.87 | 0.081 | 0.87 | 0.397 |
| AnG | 2.23 | 0.040 | 3.39 | 0.004* | 0.30 | 0.765 |
| vIPS | 1.33 | 0.204 | 1.06 | 0.304 | 0.54 | 0.600 |

One sample $t$ -tests comparing goal-relevant and goal-irrelevant feature information during memory retrieval to chance (zero) and paired sample $t$ -tests comparing goal-relevant to goal-irrelevant feature information.
\* $p < 0.00625$ following multiple comparisons correction for 8 ROIs

We first tested whether activity patterns evoked during perception and memory contained information about stimulus features (color, object). We computed the Fisher *z*-transformed Pearson’s correlation between every pair of *t*-maps from a given subject and ROI, separately for perception and memory. Note that within-stimulus identity correlations were excluded because the correlation coefficient was 1.0. We then averaged correlation values across stimulus pairs that shared a color feature (within-color correlations; e.g., blue bird - blue insect), stimulus pairs that shared an object category feature (within-object correlations; e.g., blue insect - red insect), and stimulus pairs that shared neither color nor object category (across-both correlations; e.g., red insect - yellow backpack; see Fig. 4*A*). The average across-both correlation functioned as a “baseline” and was subtracted (a) from the average within-color correlation to produce a measure of color information, and (b) from the average within-object correlation to produce a measure of object information. Thus, positive values for these measures reflected the presence of stimulus feature information. Because the perception and memory tasks did not require subjects to report the features of the stimuli (either in Experiment 1 or 2), feature information values could not be explained in terms of planned motor responses. Color and object feature information measures were computed for each subject, ROI, and run type (perception, memory). We used mixed effects ANOVAs to test whether feature information varied as a function of region (within-subject factor), run type (within-subject factor), feature dimension (within-subject factor), and/or experiment (across-subject factor). We also performed one sample *t*-tests to assess whether feature information was above chance (zero) during perception and memory.

**Figure 4.**
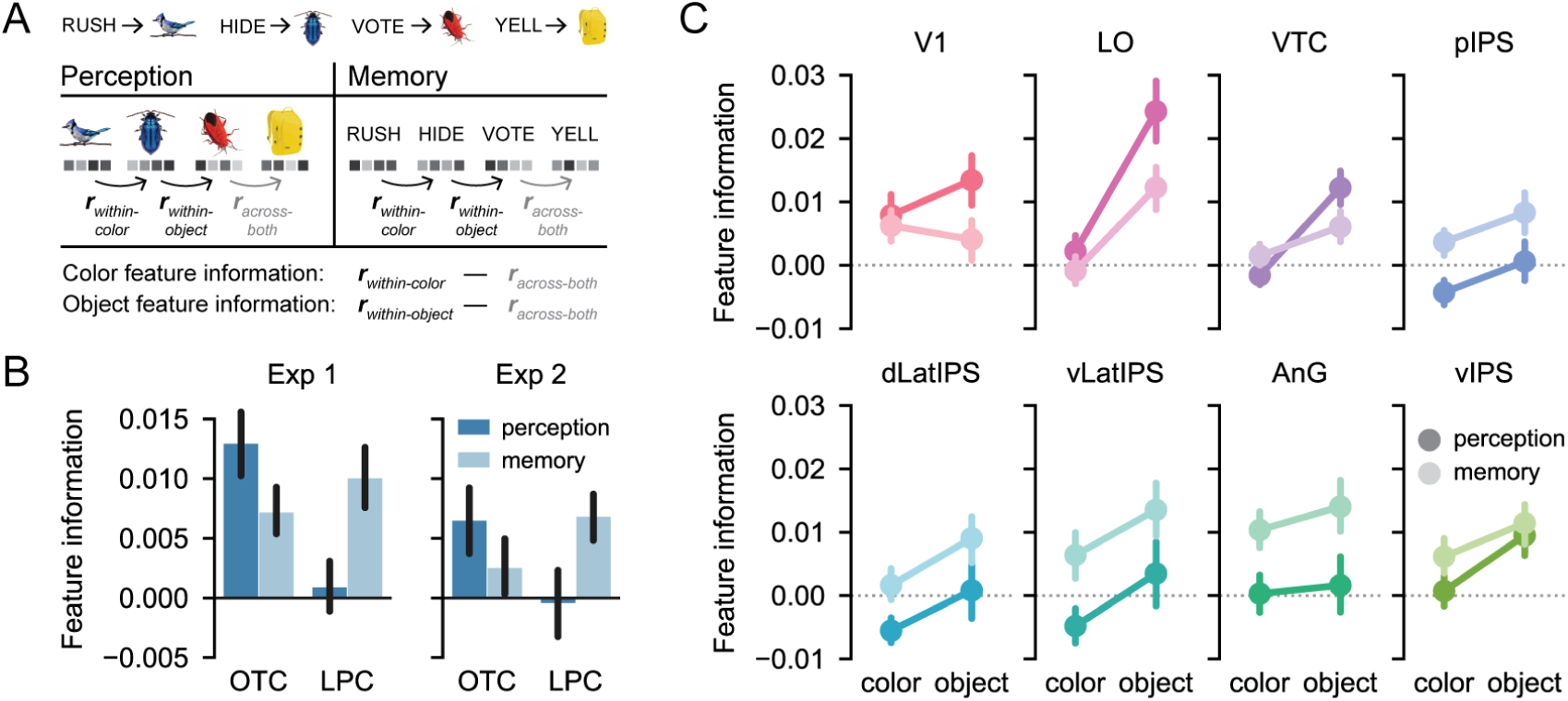
Color and object feature information during perception and memory. ***A***, Patterns of BOLD activity evoked by each stimulus were separately estimated for perception and memory runs of each experiment. We computed color and object feature information metrics for perception and memory patterns separately. To do this, we compared correlations between stimuli that shared either color or object (within-color and within-object correlations) to correlations between stimuli that shared neither feature (across-both correlations). See also Materials and Methods. ***B***, Across both experiments, the relative strength of perceptual and mnemonic feature information differed between OTC and LPC (*F*_1,32_ = 29.27, *p* < 0.001). OTC contained marginally weaker feature information during memory than during perception (*F*_1,32_ = 3.93, *p* = 0.056), while LPC contained stronger feature information during memory than during perception (*F*_1,32_ = 11.65, *p* = 0.002). Bars represent mean *±* SEM across 17 subjects. ***C***, Color and object feature information during perception and memory plotted separately for each ROI, collapsed across experiment. Points represent mean *±* SEM across 34 subjects. See Table 1 for results of one sample *t*-tests comparing feature information to chance for each ROI, run type, and feature dimension.

We next tested whether feature-level and stimulus-level information was preserved from perception to memory (reinstated). We computed the Fisher *z*-transformed Pearson’s correlation between perception and memory patterns for every pair of stimuli, separately for each subject and ROI. Excluding within-stimulus correlations, we then averaged correlation values across stimulus pairs that shared a color feature (within-color correlations; e.g., blue insect - blue bird), stimulus pairs that shared an object category feature (within-object correlations; e.g., blue insect - red insect), and stimulus pairs that shared neither color nor object category (across-both correlations; e.g., blue insect - yellow backpack; see Fig. 5*A*). The average across-both correlation functioned as a “baseline” and was subtracted (a) from the average within-color correlation to produce a measure of color reinstatement, and (b) from the average within-object correlation to produce a measure of object reinstatement. Note that these metrics are equivalent to those described in the prior section, but with correlations computed across perception and memory rather than within perception and memory. Thus, positive values for these measures reflected the preservation of feature information across perception and memory, or feature reinstatement. We used mixed effects ANOVAs to test whether feature reinstatement varied as a function of region (within-subject factor), feature dimension (within-subject factor), and/or experiment (across-subject factor). We also performed one sample *t*-tests to assess whether feature reinstatement was above chance (zero). To produce a measure of stimulus reinstatement that was comparable to our measures of feature reinstatement, we averaged within-stimulus correlation values (e.g., blue insect - blue insect) and then subtracted the same baseline (the average of across-both correlations). We performed one sample *t*-tests to assess whether stimulus reinstatement was above chance (zero). Then, we evaluated whether stimulus reinstatement could be accounted for by color and object feature reinstatement or whether it exceeded what would be expected by additive feature reinstatement. To do this we compared stimulus reinstatement to summed color and object feature reinstatement. We used mixed effects ANOVAs to test whether reinstatement varied as a function of region (within-subject factor), reinstatement level (stimulus, summed features; within-subject factor), and/or experiment (across-subject factor).

**Figure 5.**
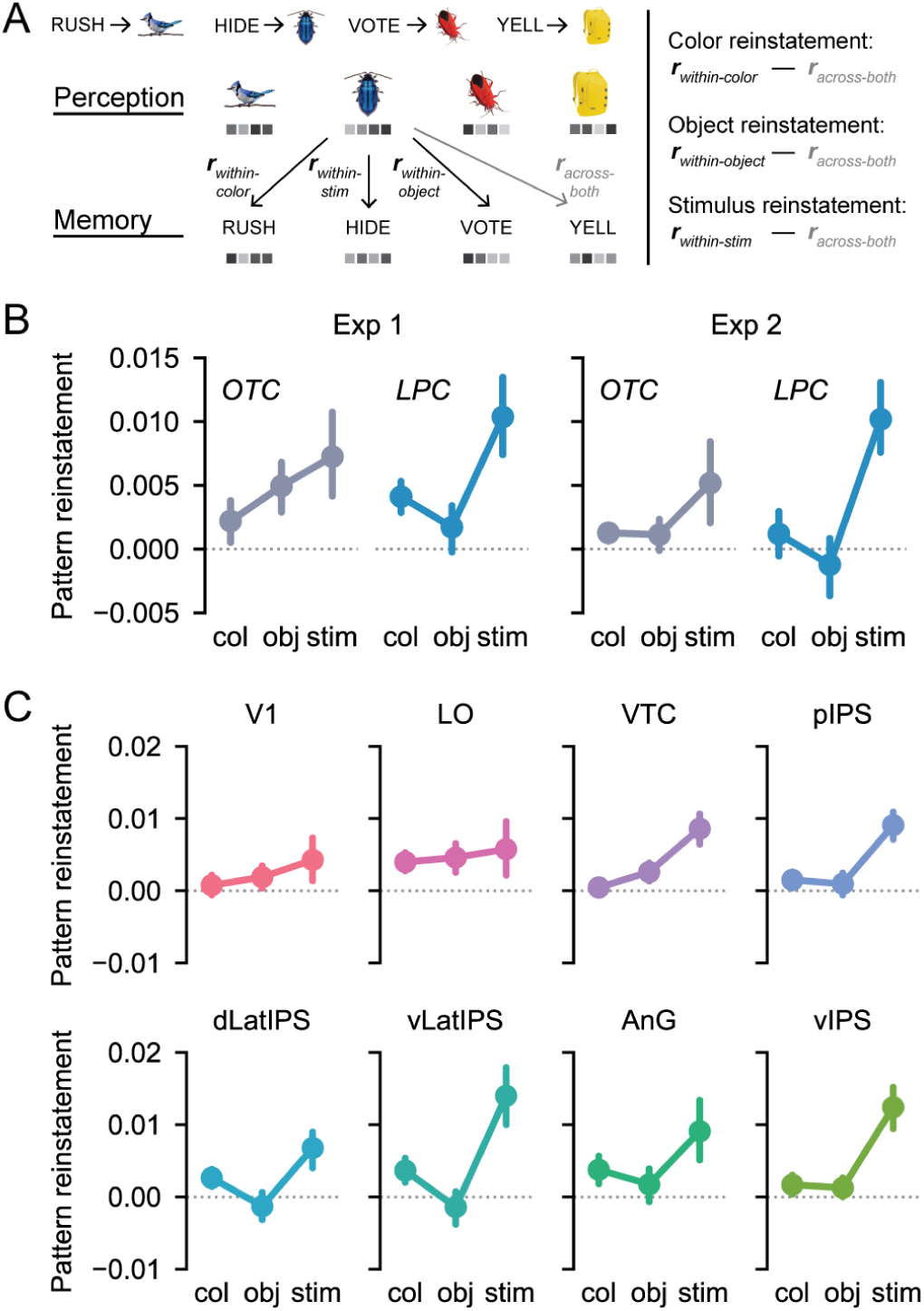
Feature and stimulus reinstatement effects. ***A***, Stimulus-evoked BOLD activity patterns were correlated across perception and memory. To quantify feature information shared across perception and memory (feature reinstatement), we compared across-stage correlations between stimuli that shared either color or object (within-color and within-object correlations) to correlations between stimuli that shared neither feature (across-both correlations). These metrics are analogous to those shown in Figure 4*A*, but were performed across stages rather than within them. To create a comparable measure of stimulus reinstatement, we compared across-stage correlations between the same stimulus (within-stimulus correlations) to correlations between stimuli that shared neither feature (across-both correlations). See also Materials and Methods. ***B***, Feature and stimulus reinstatement plotted separately for OTC and LPC and for each experiment. Across both experiments, stimulus reinstatement reliably exceeded summed levels of color and object feature reinstatement in LPC (*F*_1,32_ = 5.46, *p* = 0.026). This effect was marginally greater than the effect observed in OTC (*F*_1,32_ = 3.59, *p* = 0.067), where stimulus reinstatement was well-accounted for by summed color and object feature reinstatement (*F*_1,32_ = 0.35, *p* = m0.560). Bars represent mean *±* SEM across 17 subjects. ***C***, Color reinstatement, object reinstatement, and stimulus reinstatement plotted separately for each ROI, collapsed across experiment. Points represent mean *±* SEM across 34 subjects. See Table 3 for results of *t*-tests comparing feature reinstatement to chance for each ROI and feature dimension and comparing stimulus reinstatement to summed feature reinstatement for each ROI.

To test whether task goals influenced feature information during memory, we recomputed color and object feature information separately using *t*-maps estimated from the color and object memory tasks in Experiment 2. We averaged these feature information values into two conditions: goal-relevant (color information for the color memory task; object information for the object memory task) and goal-irrelevant (color information during the object memory task; object information during the color memory task). We used repeated measures ANOVAs to test whether feature information varied as function of region and goal-relevance (within-subjects factors). We also performed one sample *t*-tests to assess whether goal-relevant feature information and goal-irrelevant feature information were above chance (zero) during perception and memory.

## Results

### Behavior

Subjects in both experiments completed a minimum of 6 test blocks during the associative learning session prior to scanning (Exp 1: *M* = 6.65, SD = 0.79; Exp 2: *M* = 6.91, SD = 0.69). During fMRI perception runs, subjects performed the target detection task with high accuracy (Exp 1: *M* = 89.0%, SD = 6.8%; Exp 2: M = 91.6%, SD = 2.7%). In Experiment 1, subjects reported that they experienced vivid memory on a mean of 86.4% of fMRI memory trials (SD = 8.4%), weak memory on 10.4% of trials (SD = 7.1%), no memory on 1.3% of trials (SD = 1.8%), and did not respond on the remaining 1.8% of trials (SD = 2.3%). In Experiment 2, the mean percentage of vivid, weak, no memory, and no response trials was 86.1% (SD = 9.0%), 5.2% (SD = 6.1%), 3.4% (SD = 5.2%), and 5.4% (SD = 6.2%), respectively. The percentage of vivid memory responses did not significantly differ between Experiment 1 and Experiment 2 (*t*_32_ = 0.13, *p* = 0.897, independent samples *t*-test). Within each experiment, there were no differences in the percentage of vivid memory responses across stimuli with different color features (Exp 1: *F*_3,48_ = 1.19, *p* = 0.323; Exp 2: *F*_3,48_ = 0.48, *p* = 0.697; repeated measures ANOVAs) or different object features (Exp 1: *F*_7,112_ = 1.68, *p* = 0.121; Exp 2: *F*_7,112_ = 1.28, *p* = 0.266).

### Feature information during perception and memory

Across both experiments, we quantified the strength of color and object feature information contained in stimulus-evoked BOLD patterns throughout OTC and LPC. We did this separately for patterns evoked by perception and memory (Fig. 4*A*; see Materials and Methods). Of critical interest was whether the relative strength of perceptual and mnemonic feature information differed across LPC and OTC (Fig. 3). We entered feature information values from all ROIs into an ANOVA with factors of ROI group (OTC, LPC), run type (perception, memory), feature dimension (color, object), and experiment (Exp 1, Exp 2). Critically, the relative strength of perception and memory-based feature information differed across LPC and OTC, as reflected by a highly significant interaction between ROI group and run type (*F*_1,32_ = 29.27, *p* < 0.001; Fig. 4*B*). This effect did not differ across experiments (ROI group x run type x experiment interaction: *F*_1,32_ = 0.55, *p* = 0.462; Fig. 4*B*).

In LPC, feature information was reliably stronger during memory than during perception (main effect of run type: *F*_1,32_ = 11.65, *p* = 0.002; Fig. 4*B*), with no difference in this effect across individual LPC ROIs (run type x ROI interaction *F*_4,128_ = 1.55, *p* = 0.192; Fig. 4*C*). Averaging across the color and object dimensions, and also experiments, feature information was above chance during memory (*t*_33_ = 4.79, *p* < 0.001; one sample *t*-test), but not during perception (*t*_33_ = 0.14, *p* = 0.892; Fig. 4*B*). In Table 1 we report the results of one sample *t*-tests comparing feature information to chance for each LPC ROI, run type, and feature dimension separately. There was also a marginally significant main effect of feature dimension in LPC (*F*_1,32_ = 3.95, *p* = 0.056), with somewhat stronger object information than color information. This marginally significant main effect of feature dimension did not interact with run type (*F*_1,32_ = 0.004, *p* = 0.952).

In OTC, we observed a pattern opposite to LPC: feature information was marginally stronger during perception than during memory (main effect of run type: *F*_1,32_ = 3.93, *p* = 0.056; Fig. 4*B*). Again, this effect did not differ across individual OTC ROIs (run type x ROI interaction: *F*_2,64_ = 1.72, *p* = 0.187; Fig. 4*C*). Averaging across the color and object dimensions, and also experiments, feature information was above chance both during perception (*t*_33_ = 4.68, *p* < 0.001) and during memory (*t*_33_ = 3.01, *p* = 0.005; Fig. 4*B*). Table 1 includes comparisons of feature information to chance for each OTC ROI, run type, and feature dimension. There was a significant main effect of feature dimension in OTC (*F*_1,32_ = 18.59, *p* < 0.001), with stronger object information than color information. This main effect of feature dimension interacted with run type (*F*_1,32_ = 4.90, *p* = 0.034), reflecting a relatively stronger difference between color and object information during perception than during memory. Together, these results indicate that feature information was differentially expressed in LPC and OTC depending on whether the stimulus was internally generated (memory retrieval) or external (perception).

### Feature-level and stimulus-level reinstatement

We next quantified reinstatement of feature information. Whereas the prior analyses separately quantified feature information during perception and memory retrieval, here we tested whether feature-specific activity patterns were preserved across perception and memory (Fig. 5*A*; see Materials and Methods). Because perception and memory trials had no overlapping visual elements, any feature information preserved across stages must reflect memory retrieval of stimulus features. We entered feature reinstatement values from all ROIs into an ANOVA with factors of ROI group (OTC, LPC), feature dimension (color, object), and experiment (Exp 1, Exp 2). There was no reliable difference in the strength of feature reinstatement between OTC and LPC (main effect of ROI group: *F*_1,32_ = 0.90, *p* = 0.350). There was also no overall difference in the magnitude of color versus object reinstatement (main effect of feature dimension: *F*_1,32_ = 0.82, *p* = 0.373), and this effect did not vary between OTC and LPC (feature dimension x ROI group interaction: *F*_1,32_ = 2.41, *p* = 0.130). There was a marginal main effect of experiment on feature reinstatement (*F*_1,32_ = 3.10, *p* = 0.088), with numerically lower feature reinstatement in Experiment 2 (where subjects recalled only one feature) than in Experiment 1 (where subjects recalled the entire image) (Fig. 5*B*). When collapsing across color and object dimensions, feature reinstatement was above chance in OTC in both Experiment 1 (*t*_16_ = 2.37, *p* = 0.031; one sample *t*-test) and Experiment 2 (*t*_16_ = 2.33, *p* = 0.033). In LPC, feature reinstatement was above chance in Experiment 1 (*t*_16_ = 2.58, *p* = 0.020), but not in Experiment 2 (*t*_16_ = -0.007, *p* = 0.995). Thus, the task demands in Experiment 2 may have had a particular influence on LPC feature representations–a point we examine in the next section. In Table 2 we report the results of one sample *t*-tests comparing feature reinstatement to chance for each ROI and feature dimension (see also Fig. 5*C*).

We next tested whether stimulus-level information was reinstated. As a first step analysis, we tested whether within-stimulus correlations (i.e., perception and memory trials corresponding to the same stimulus) were greater than across-both correlations (i.e., perception and memory trials corresponding to stimuli with no common features). Indeed, within-stimulus correlation values were significantly greater than across-both correlations in LPC (*t*_33_ = 5.03, *p* < 0.001; paired sample *t*-test) and OTC (*t*_33_ = 2.71, *p* = 0.011; Fig. 5*B*). We refer to this difference between within-stimulus and across-both correlations as a measure of stimulus reinstatement (Fig. 5*A*). Importantly, the fact that we observed positive stimulus reinstatement values in LPC and OTC could be entirely driven by reinstatement of one or both features (color and/or object). To test whether stimulus reinstatement could be fully accounted for by reinstatement of object and color information, we compared stimulus reinstatement values (within-stimulus correlations minus across-both correlations; Fig. 5*A*) against a summed measure of color and object reinstatement values. Reinstatement values from all ROIs were entered into an ANOVA with factors of ROI group (OTC, LPC), reinstatement level (stimulus reinstatement, summed features), and experiment (Exp 1, Exp 2). There was a significant main effect of reinstatement level (*F*_1,32_ = 4.31, *p* = 0.046), with stimulus reinstatement larger than summed feature reinstatement. There was a marginally significant difference in the magnitude of this effect between OTC and LPC (reinstatement level interaction x ROI group: *F*_1,32_ = 3.59, *p* = 0.067; Fig. 5*B*). In LPC, stimulus reinstatement reliably exceeded summed feature reinstatement (main effect of reinstatement level: *F*_1,32_ = 5.46, *p* = 0.026; Fig. 5*B*). This effect did not differ across experiments (reinstatement level x experiment interaction: *F*_1,32_ = 0.81, *p* = 0.375) or across LPC ROIs (reinstatement level x ROI interaction: *F*_4,128_ = 0.95, *p* = 0.438; Fig. 5*C*). In Table 2 we report the results of paired *t*-tests comparing stimulus reinstatement to summed feature reinstatement for each LPC ROI. In OTC, stimulus reinstatement did not significantly differ from summed feature reinstatement (main effect of reinstatement level: *F*_1,32_ = 0.35, *p* = 0.560; Fig. 5*B*), with no difference across experiments (reinstatement level x experiment interaction: *F*_1,32_ = 0.30, *p* = 0.590) and a marginal difference across ROIs (reinstatement level x ROI interaction: *F*_2,64_ = 2.58, *p* = 0.084). Tests in individual OTC ROIs (Table 2) showed that stimulus reinstatement significantly exceeded summed feature reinstatement in VTC only. These results replicate prior evidence of stimulus-specific reinstatement in LPC (Kuhl and Chun, 2014; Lee and Kuhl, 2016; Xiao et al., 2017) and VTC (Lee et al., 2012), but provide unique insight into the relative strength of feature-vs. stimulus-specific information across these regions.

### Goal-dependence of mnemonic feature information

In a final set of analyses, we tested whether feature information expressed in LPC during memory retrieval was influenced by goals. Using data from Experiment 2 only, we recomputed color and object feature information separately for trials where recalling the color of the stimulus was relevant and trials where recalling the object category of the stimulus was relevant (see Materials and Methods). Of interest was the comparison between goal-relevant feature information (e.g., color information on color memory trials) and goal-irrelevant feature information (e.g., color information on object memory trials). Because there is a strong body of evidence suggesting that dorsal and ventral parietal regions are differentially sensitive to top-down vs. bottom-up visual attention (Corbetta and Shulman, 2002), for this analysis we specifically tested whether sensitivity to retrieval goals varied across dorsal and ventral LPC subregions (Fig. 6*A*; see Materials and Methods).

**Figure 6.**
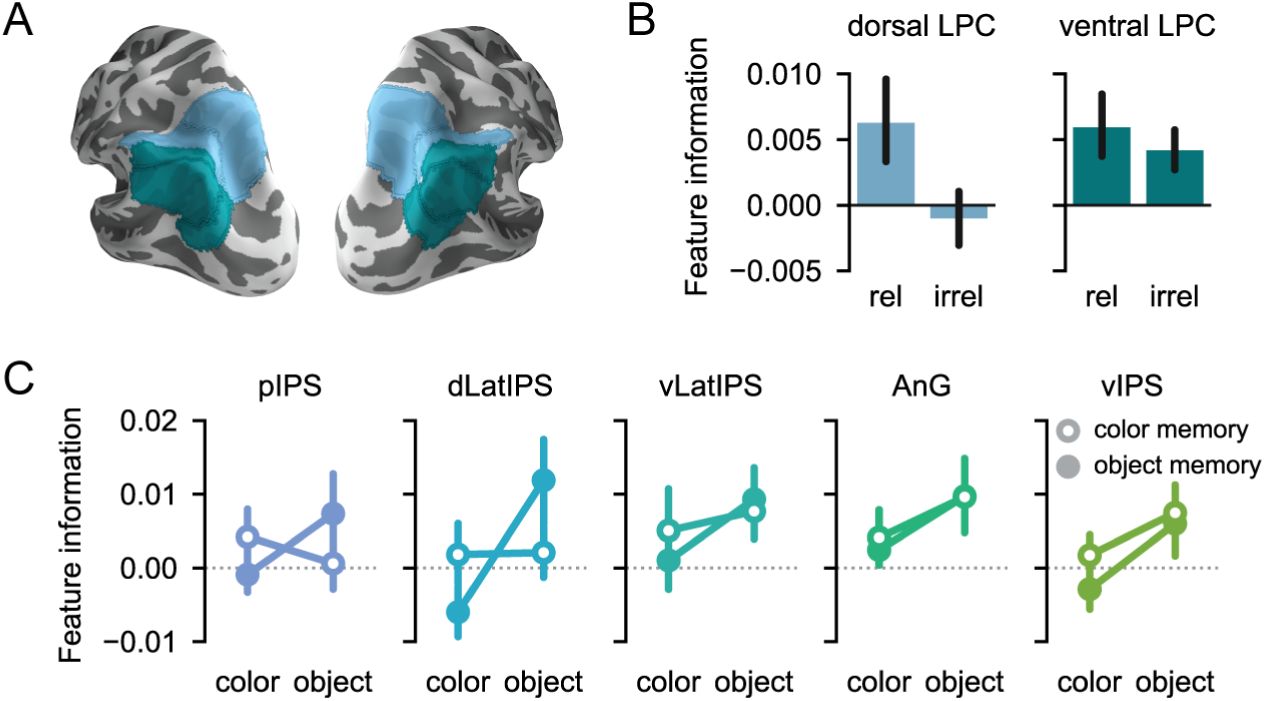
Feature information during memory as a function of goal-relevance. *A*, ROIs from Figure 3 grouped according to a dorsal/ventral division along the intraparietal sulcus (see Materials and Methods). *B*, The effect of goal-relevance on mnemonic feature information differed significantly between dorsal and ventral LPC subregions (*F*_1,16_ = 9.05, *p* = 0.008). In dorsal LPC, goal-relevant feature information was stronger than goal-irrelevant feature information (*F*_1,16_ = 5.30, *p* = 0.035). In ventral LPC, there was no effect of goal-relevance on feature information (*F*_1,16_ = 0.61, *p* = 0.447), and both goal-relevant (*t*_16_ = 2.48, *p* = 0.025) and goal-irrelevant (*t*_16_ = 2.64, *p* = 0.018) feature information were represented above chance. Bars represent mean *±* SEM across 17 subjects. *C*, Color and object feature information during color and object memory tasks plotted separately for each LPC ROI. Points represent mean *±* SEM across 17 subjects. See Table 3 for results of *t*-tests assessing mnemonic feature information according to goal-relevance for each ROI.

To first broadly compare LPC and OTC, we entered memory-based feature information values from all ROIs into an ANOVA with factors of ROI group (OTC, LPC) and goal-relevance (relevant, irrelevant). The relative strength of goal-relevant vs. goal-irrelevant feature information differed across LPC and OTC, as revealed by a significant ROI group x goal-relevance interaction (*F*_1,16_ = 7.30, *p* = 0.016). In LPC, goal-relevant feature information was numerically, but not significantly, greater than goal-irrelevant feature information (main effect of goal-relevance: *F*_1,16_ = 2.53, *p* = 0.131), whereas in OTC the pattern was qualitatively opposite (*F*_1,16_ = 1.51, *p* = 0.237).

Next, to test the more targeted question of whether goal sensitivity varied between dorsal and ventral LPC subregions, we entered memory-based feature information values from LPC ROIs into an ANOVA with factors of LPC subregion (dorsal LPC, ventral LPC) and goal-relevance (relevant, irrelevant). In line with our hypothesis, there was a robust interaction between LPC subregion and goal-relevance (*F*_1,16_ = 9.05, *p* = 0.008; Fig. 6*B*). Namely, there was reliably stronger goal-relevant than goal-irrelevant feature information in dorsal LPC (main effect of goal-relevance: *F*_1,16_ = 5.30, *p* = 0.035). This effect did not differ across individual dorsal LPC ROIs (goal-relevance x ROI interaction: *F*_1,16_ = 1.01, *p* = 0.330; Fig. 6*C*). In dorsal LPC, goal-relevant feature information marginally exceeded chance (goal-relevant: *t*_16_ = 1.93, *p* = 0.072; one sample *t*-test) whereas goal-irrelevant feature information did not differ from chance (*t*_16_ = -0.49, *p* = 0.628; Fig. 6*B*). In contrast to the pattern observed in dorsal LPC, feature information was not significantly influenced by goals in ventral LPC (main effect of goal-relevance: *F*_1,16_ = 0.61, *p* = 0.447), nor did this effect vary across ventral LPC ROIs (goal-relevance x ROI interaction: *F*_2,32_ = 0.16, *p* = 0.855). In fact, both goal-relevant and goal-irrelevant information were significantly above chance in ventral LPC (goal-relevant: *t*_16_ = 2.48, *p* = 0.025; goal-irrelevant: *t*_16_ = 2.64, *p* = 0.018; Fig. 6*B*). Thus, the interaction between dorsal vs. ventral LPC and goal-relevance was driven primarily by a difference in the strength of goal-irrelevant feature information. Indeed, goal-irrelevant feature information was significantly stronger in ventral LPC than in dorsal LPC (*t*_16_ = 3.15, *p* = 0.006; paired sample *t*-test; Fig. 6*B*), whereas the strength of goal-relevant feature information did not significantly differ across ventral and dorsal LPC (*t*_16_ = -0.19, *p* = 0.850). In Table 3 we report the results of one sample *t*-tests comparing goal-relevant and goal-irrelevant feature information to chance as well as paired sample *t*-tests comparing goal-relevant to goal-irrelevant feature information, separately for each LPC and OTC ROI (see also Fig. 6*C*). Collectively, these findings provide novel evidence for a functional distinction between memory representations in dorsal and ventral LPC, with top-down memory goals biasing feature representations toward relevant information in dorsal LPC, but not ventral LPC. Importantly, because there was no evidence of preferential representation of goal-relevant feature information during memory retrieval in OTC, the bias observed in dorsal LPC was not inherited from earlier visual regions.

## Discussion

Here, across two fMRI experiments, we show that OTC and LPC differentially represent stimulus features during perception and memory retrieval. In OTC, color and object feature information were stronger during perception than during memory, whereas in LPC, feature information was stronger during memory than during perception. Despite these biases, we observed that stimulus-specific patterns evoked in LPC during perception were reinstated during memory retrieval. In Experiment 2 we found that top-down retrieval goals biased dorsal LPC representations toward relevant features, whereas ventral LPC represented both relevant and irrelevant features. Collectively, these findings reveal that LPC plays a role in representing feature- and stimulus-specific information during memory retrieval and in adapting these representations to satisfy current goals.

### Transformation of representations from OTC to LPC

Traditionally, cortical memory reactivation has been studied in sensory regions, with the idea being that memory-related activity patterns are a degraded copy of earlier perceptual activity patterns (O’Craven and Kanwisher, 2000; Wheeler et al., 2000; Slotnick et al., 2005; Pearson et al., 2015). While this idea accounts for our results in OTC, it does not explain our results in LPC, where stimulus feature information was stronger during memory retrieval than perception. What accounts for this reversal in LPC? Notably, in ventral LPC, univariate BOLD responses tend to be higher during successful memory retrieval than during perception (Daselaar, 2009; Kim et al., 2010). These univariate increases may be driven by strong anatomical (Cavada and Goldman-Rakic, 1989; Clower et al., 2001) and functional (Vincent et al., 2006; Kahn et al., 2008) connections from medial temporal lobe (MTL) regions that are critical for recollection. Similarly, stronger pattern-based information in ventral LPC during retrieval may reflect this same intrinsic drive from the MTL. Interestingly, recent evidence from rodents also indicates that parietal cortex is biased towards memory-based representations (Akrami et al., 2018). Namely, neurons in rat posterior parietal cortex were shown to carry more information about sensory stimuli from prior trials than from the current trial, even when prior trials were not behaviorally relevant. While it thus seems likely that at least some regions of LPC (e.g., ventral LPC) are intrinsically biased toward memory representations, the fact that we also observed this same bias in dorsal LPC is somewhat more surprising given that dorsal LPC includes several retinotopically-organized regions (Silver and Kastner, 2009) that receive strong input from early visual areas (Lewis and Van Essen, 2000). However, given that dorsal LPC is thought to be involved in visual attention (Corbetta and Shulman, 2002), it is possible that the bias towards memory representations in this region can be attributed to differences in attentional demands across our perception and memory tasks. Namely, if our memory task was more attentionally demanding than our perception task–which is plausible, but difficult to test–then feature representations in dorsal LPC may have been amplified by attention during memory retrieval (Silver, 2005; Sprague and Serences, 2013; Ester et al., 2016). In either case, the fact that dorsal LPC actively represents visual features of retrieved stimuli is an important finding.

Recently, Xiao et al. (2017) reported a finding complementary to ours. specifically, they found that stimulus-specific representations were relatively stronger in frontoparietal regions during retrieval than perception, arguing that representations in OTC during perception are “transformed” to frontoparietal regions during retrieval. However, as Xiao and colleagues note, they did not measure feature-level representations and the apparent transformation of stimulus-level representations from OTC to frontoparietal cortex may reflect a difference in the features each region represents. Thus, our findings support theirs but also constitute an important advance by showing that the same stimulus features represented in OTC during perception are re-represented in LPC during retrieval. Moreover, and as we detail in the final section, considering feature-level information is essential for testing whether LPC representations of retrieved memories are flexibly aligned with goals.

### Pattern reinstatement within regions

Consistent with prior studies, we observed stimulus-specific reinstatement of perceptual patterns during memory retrieval in LPC (Buchsbaum et al., 2012; Kuhl and Chun, 2014; Ester et al., 2015; Chen et al., 2016; Lee and Kuhl, 2016; Xiao et al., 2017) and VTC (Lee et al., 2012). Interestingly, we observed reinstatement in LPC and VTC despite the fact that these regions each had a bias toward either mnemonic (LPC) or perceptual (VTC) information. While these findings may seem contradictory, it is important to emphasize that the biases we observed were not absolute. Rather, there was significant feature information in OTC during memory retrieval, and though we did not observe significant feature information in LPC during perception, other studies have reported LPC representations of perceptual stimuli (Bracci et al., 2017; Lee et al., 2017). Thus, we think it is likely that the reinstatement effects that we and others have observed co-occur with a large but incomplete transformation of stimulus representation from OTC during perception to LPC during retrieval.

Notably, the stimulus reinstatement effects that we observed in LPC could not be explained by additive reinstatement of color and object information. Likewise, our pre-scan associative learning task required subjects to learn more than just color-object feature conjunctions. Thus, LPC representations, like subjects’ memories for the stimuli, likely reflected the conjunction of more than just color and object information. This is consistent with theoretical arguments and empirical evidence suggesting that parietal cortex–and, in particular, angular gyrus–serves as a multimodal hub that integrates event features in memory (Shimamura, 2011; Wagner et al., 2015; Bonnici et al., 2016; Yazar et al., 2017). Given that ventral LPC is frequently implicated in semantic processing (Binder and Desai, 2011), it is likely that stimulus-specific representations in ventral LPC reflect a combination of perceptual and semantic information. In contrast, stimulus-specific representations in dorsal LPC and VTC, which are components of two major visual pathways, potentially reflect combinations of high-level but fundamentally perceptual features.

### influence of retrieval goals on LPC representations

Substantial evidence from electrophysiological (Toth and Assad, 2002; Freedman and Assad, 2006; Ibos and Freedman, 2014) and BOLD (Liu et al., 2011; Erez and Duncan, 2015; Bracci et al., 2017; Vaziri-Pashkam and Xu, 2017; Long and Kuhl, 2018) measurements indicates that LPC representations of perceptual events are influenced by top-down goals. Our results provide novel evidence that, in dorsal LPC, specific features of a remembered stimulus are dynamically strengthened or weakened according to the current goal. This finding provides a critical bridge between perception-based studies emphasizing the role of LPC in goal-modulated stimulus coding and memory-based studies that have found representations of remembered stimuli in LPC. Importantly, because we did not require subjects to behaviorally report any feature information, the representations we observed cannot be explained in terms of action planning (Andersen and Cui, 2009). However, the fact that we observed goal-modulated feature coding in dorsal, but not ventral, LPC is consistent with theoretical accounts arguing that dorsal LPC is more involved in top-down attention whereas ventral LPC is more involved in bottom-up attention (Corbetta and Shulman, 2002). Cabeza et al. (2008) has argued that LPC’s role in memory can similarly be explained in terms of top-down and bottom-up attentional processes segregated across dorsal and ventral LPC. However, from this account, LPC is not thought to actively represent mnemonic content. Thus, while our findings support the idea that dorsal and ventral LPC are differentially involved in top-down vs. bottom-up memory processes, they provide critical evidence that these processes involve active representation of stimulus features.

Interestingly, although we observed no difference between goal-relevant and goal-irrelevant feature information in ventral LPC, both were above chance. This is consistent with the idea that ventral LPC represents information received from the MTL, perhaps functioning as an initial mnemonic buffer (Baddeley, 2000; Vilberg and Rugg, 2008; Kuhl and Chun, 2014; Sestieri et al., 2017). Ventral LPC representations may then be selectively gated according to current behavioral goals, with goal-relevant information propagating to dorsal LPC. This proposal is largely consistent with a recent theoretical argument made by Sestieri et al. (2017). However, it differs in the specific assignment of functions to LPC subregions. Whereas Sestieri et al. (2017) argue that dorsal LPC is contributing to goal-directed processing of perceptual information only, our results indicate that dorsal LPC also represents mnemonic information according to current goals. Given the paucity of experiments examining the influence of goals on mnemonic representations in LPC (c.f. Kuhl et al., 2013), additional evidence is needed. However, our findings provide important evidence, motivated by existing theoretical accounts, that goals differentially influence feature representations across LPC subregions.

### Conclusions

In summary, we show that LPC not only actively represents features of remembered stimuli, but that these LPC feature representations are stronger during memory retrieval than perception. Moreover, whereas ventral LPC automatically represents stimulus features irrespective of goals, dorsal LPC feature representations are flexibly and dynamically influenced to match top-down goals. Collectively, these findings provide novel insight into the functional significance of memory representations in LPC.

## Acknowledgements

This work was made possible by generous support from the Lewis Family Endowment to the University of Oregon Robert and Beverly Lewis Center for Neuroimaging. S.E.F was supported by the NSF Graduate Research Fellowship Program (DGE-1342536) and the NEI Visual Neuroscience Training Program (T32-EY007136). B.A.K. was supported by an NSF CAREER Award (BCS-1752921).

